# Gpufit: An open-source toolkit for GPU-accelerated curve fitting

**DOI:** 10.1101/174110

**Authors:** Adrian Przybylski, Björn Thiel, Jan Keller-Findeisen, Bernd Stock, Mark Bates

## Abstract

We present a general purpose, open-source software library for estimation of non-linear parameters by the Levenberg-Marquardt algorithm. The software, Gpufit, runs on a Graphics Processing Unit (GPU) and executes computations in parallel, resulting in a significant gain in performance. We measured a speed increase of up to 42 times when comparing Gpufit with an identical CPU-based algorithm, with no loss of precision or accuracy. Gpufit is designed such that it is easily incorporated into existing applications or adapted for new ones. Multiple software interfaces, including to C, Python, and Matlab, ensure that Gpufit is accessible from most programming environments. The full source code is published as an open source software repository, making its function transparent to the user and facilitating future improvements and extensions. As a demonstration, we used Gpufit to accelerate an existing scientific image analysis package, yielding significantly improved processing times for super-resolution fluorescence microscopy datasets.

## Introduction

Optimization algorithms are widely used in science and engineering. In particular, when comparing data with a model function, numerical optimization methods may be applied to establish the suitability of the model and to determine the parameters which best describe the observations. One such method, which is generally applicable to models which depend non-linearly on a set of parameters, is the Levenberg-Marquardt algorithm (LMA),^1^ and this has become a standard approach for non-linear least squares curve fitting.^2,3^

Although the LMA is, in itself, an efficient algorithm, applications requiring many iterations of this procedure may encounter limitations due to the sheer number of calculations involved. The time required for the convergence of a fit, or a set of fits, can determine an application’s feasibility, e.g. in the context of real-time data processing and feedback systems. Alternatively, in the case of very large datasets, the time required to solve a particular optimization problem may prove impractical.

In recent years, advanced graphics processing units (GPUs) and the development of general purpose GPU programming have enabled fast and parallelized computing by shifting calculations from the CPU to the GPU.^4^ The large number of independent computing units available on modern GPUs enables the rapid execution of many instructions in parallel, with an overall computation power far exceeding that of a CPU. Languages such as CUDA C and OpenCL allow GPU-based programs to be developed in a manner similar to conventional software, but with an inherently parallelized structure. These developments have led to the creation of new GPU-accelerated tools, such as the MAGMA linear algebra library,^5^ for example.

Here, we present Gpufit: a GPU-accelerated implementation of the Levenberg-Marquardt algorithm. Gpufit was developed to meet the need for a high performance, general-purpose nonlinear curve fitting library which is publicly available and open source. As expected, this software exhibits significantly faster execution than the equivalent CPU-based code, with no loss of precision or accuracy. In this report we discuss the design of Gpufit, characterize its performance in comparison to other CPU-based and GPU-based algorithms, and demonstrate its use in a scientific data analysis application.

## Results

### Software design

The Gpufit library was designed to meet several criteria: *(i)* the software should make efficient use of the GPU resources in order to maximize execution speed, *(ii)* the interface should not require detailed knowledge of the GPU hardware, *(iii)* the source code should be modular and extendable, and *(iv)* the software should be accessible from multiple programming environments.

Gpufit implements the LMA entirely on the GPU, minimizing data transfers between CPU and GPU memory. Copying memory between the CPU and the GPU is a slow operation, and it is most efficient to handle large blocks of data at once. Therefore, Gpufit is designed to process multiple fits simultaneously, each with the same model function and data size, but allowing for unique starting parameters for each fit. Large tasks (e.g. large numbers of fits) are divided into chunks, in order to balance processing and data transfer times. At the start of a calculation, a chunk of input data is copied to the GPU global memory, and upon completion the results are transferred back to CPU memory.

GPU architecture is based on a set of parallel multiprocessors, which divide computations between blocks of processing threads, as illustrated in Supplementary Fig. S1. The efficiency of a GPU-based program depends on how these computing resources are used. While determining how best to parallelize the LMA, we found that different parts of the algorithm were most efficiently implemented with different parallelization strategies. For example, point-wise operations such as computation of the model function and its partial derivatives were more efficiently parallelized along the data coordinate index, meaning that each thread computes one model and derivative value at a particular coordinate. Other steps, such as the calculation of the Hessian matrix, were more efficiently parallelized along the index of the matrix element, i.e. with each thread assigned to calculate one element of the matrix. To accommodate the necessity for parallelizing different parts of the LMA in different ways, we structured Gpufit as a set of independent CUDA kernels, each responsible for a section of the algorithm. In this way, the blocks and threads of the GPU multiprocessors could be optimally allocated at each step. The details of the various parallelization schemes are documented in the Gpufit source code (see Code Availability).

Another design consideration was how to distribute fits across the thread blocks of the GPU. Initially, we considered calculating one fit per thread block, and one data point per thread. However, this approach yielded poor results for small fit data sizes, due to the limited numbers of threads executed on each multiprocessor. Hence, the software was modified to allow the calculation of multiple fits in each thread block or, if necessary, to spread the calculation of a single fit over multiple thread blocks. Thus, the number of threads per block was maximized, yielding improved processing speeds without imposing restrictions on the size of each fit. Finally, we note that GPU shared memory (see Fig. S1) was used for the calculation of intermediate values requiring many reads and writes (e.g. the summation of chi-square), in order to take advantage of the faster performance of shared memory blocks.

The computing resources of a GPU may vary significantly, depending on the details of the hardware. One of our aims was to avoid the necessity for any hardware-specific configuration parameters in the Gpufit interface. Therefore, Gpufit was designed to read the properties of the GPU at run-time, and automatically set parameters such as the number of blocks and threads available, and the number of fits to calculate simultaneously. This makes the use of a GPU transparent to the programmer, and allows the interface to be no different from that for a conventional curve fitting function. Moreover, it ensures that applications using Gpufit will perform optimally regardless of the machine on which they are running.

The Gpufit source code is modular, such that fit functions and goodness-of-fit estimators are separate from the core sections of the code, and new functions or estimators may be added simply (see Supplementary Information). In its initial release, Gpufit includes two different fit estimators: the standard weighted least-squares estimator (LSE), and a maximum likelihood estimator (MLE) which provides better fit results when the input data is characterized by Poisson statistics.^6^ The modular concept is illustrated schematically in Supplementary Fig. S2. This modularity allows Gpufit to be quickly adapted to new applications, or modified to accommodate future developments.

Finally, Gpufit is written in C, CUDA C, and C++, and compiles to a Dynamic Link Library (DLL) providing a C interface, making it straightforward to call Gpufit from most programming environments. Furthermore, Gpufit bindings for Matlab and Python (e.g. the pyGpufit module) were implemented, forming part of the Gpufit distribution.

### Performance characterization

We tested Gpufit by using the software to process randomized simulated datasets. The precision and accuracy of the fit results, as well as the number of fit iterations and the execution time, were measured. The input data consisted of 2D Gaussian functions defined by five parameters (see Supporting Information). Random noise (Gaussian or Poisson) was added to each data point. Test data was generated in Matlab and passed to the fit routines via their Matlab interfaces.

### Algorithm validation

An initial step in characterizing Gpufit was to verify the correct operation of the algorithm. For this purpose, we tested Gpufit against the well-established MINPACK library,^7^ evaluating the precision and accuracy of the fit results, and the convergence of the fit. Gaussian noise was added to the input data such that the signal to noise ratio (SNR) could be defined. Upon testing, the two packages yielded almost identical fit precision over a wide range of SNR values, and converged in a similar number of steps (Supplementary Fig. S3). At very high SNR, differences appear due to the limited numerical precision of floating point operations in CUDA (single precision) vs. MINPACK (double precision). Fit accuracy was checked by comparing the distributions of the fit parameters (Supplementary Fig. S4), and no systematic deviation between the output of Gpufit and MINPACK was detected. Together, these measurements demonstrate that Gpufit yields precise and accurate fit results, and converges similarly when compared to existing optimization software.

### Execution speed and fit precision

The parallel architecture of the GPU enables significant speed improvements when a computation is amenable to being divided among multiple processors. To fairly evaluate the speed improvement obtained by shifting processing tasks from the CPU to the GPU, equivalent implementations of the LMA, running on both architectures, were required. We created a library called Cpufit to serve as the reference CPU-based fitting algorithm for testing purposes (see Methods). Cpufit implements precisely the same algorithm, section by section, as Gpufit. We verified that Cpufit and Gpufit returned numerically identical fit results given identical input data.

A comparison of execution speeds, plotted against the number of fits per function call (*N*), is shown in Fig. 1. As expected, the fitting speed on the CPU remains constant as *N* changes, due to the sequential execution of each fit calculation. On the other hand, for Gpufit the speed increases with increasing *N*. The more fits that are calculated in one execution, the greater the benefit of parallelization. The crossover point at which use of the GPU becomes advantageous is, in this case, approximately *N* = 130 fits. For very large *N* the speed of GPU processing saturates, indicating that GPU resources are fully utilized. At *N* = 10^8^ fits per function call, we measured a processing speed of more than 4.5 million fits per second, approximately 42 times faster than the same algorithm executed on the CPU.

**Figure 1:**
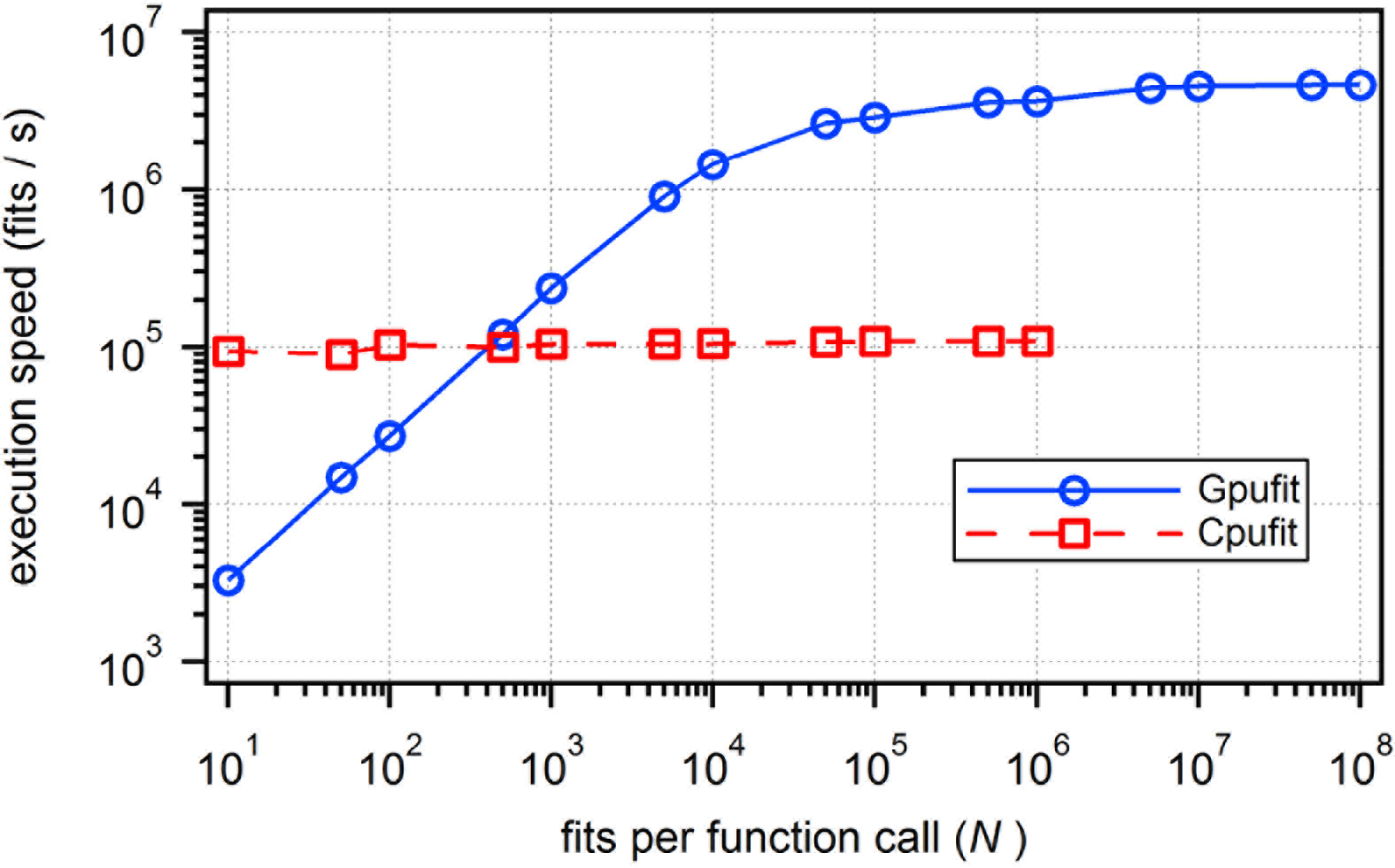
Execution speed of the Levenberg-Marquardt algorithm on the CPU or the GPU,as a function of the number of fits processed. For small numbers of fits, GPU-based curve fitting is slower than CPU-based fitting, due to the extra time spent copying data between the CPU and the GPU. For larger numbers of fits, however, the GPU significantly out-performs the CPU, due to the GPU’s parallel architecture. In this example, the maximum speed achieved was 4.65 × 10^6^ fits per second for GPU-based curve fitting (dependent on the specifics of the hardware, the fit function, and the fit data). The simulated data consisted of 2D Gaussian peaks with a size of 5 × 5 points. This data was fit to a symmetric 2D Gaussian function using the unweighted least squares estimator. All calculations were run using an NVIDIA GeForce GTX 1080 GPU and an Intel Core i7 5820K CPU (3.3 GHz). For details of the simulated data and fit parameters, see the Supplementary Information.

Measurements of the execution time for each sub-section of the Cpufit and Gpufit code reveal the bottlenecks of the CPU-based algorithm, and how the computational workload is re-distributed on the GPU. For a set of 5×10^6^ fits we measured the time duration of each step of the fitting process, and these results are shown in Fig. 2. For reference, program flowcharts for Cpufit and Gpufit, color coded according to processing time, are shown in Supplementary Fig. S5 and S6. In our tests, the most time-consuming task for Cpufit was the calculation of the model function and its derivatives, requiring more than one third of the total execution time. Gpufit completed the same calculation more than 100 times faster, due to the parallelization of the work. Similarly, all other steps in the fit process ran 10 - 100 times faster on the GPU. Gpufit includes additional operations, such as data transfers between CPU and GPU memory, however these did not impact performance when sufficient numbers of fits were processed.

**Figure 2:**
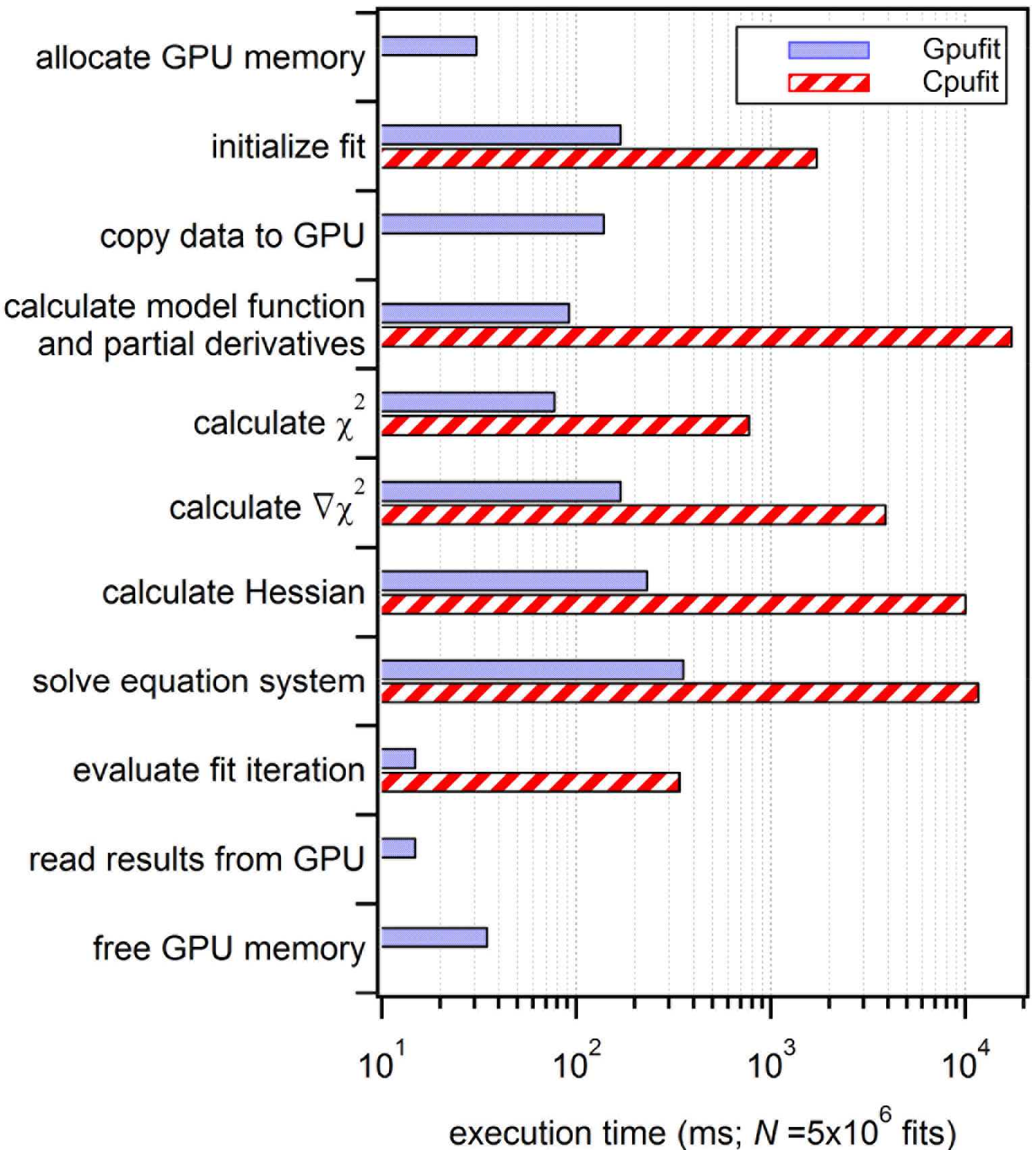
Comparison of execution times for each section of the Cpufit and Gpufit programs. The horizontal bars represent the amount of time spent in each section of the algorithm while processing a dataset consisting of *N* = 5.0×10^6^ 2D Gaussian fits. Four additional sections are required for Gpufit: GPU memory allocation, copying data to the GPU, copying results from the GPU, and GPU memory de-allocation. All sections of the fit algorithm required less time when executed on the GPU, with the largest differences corresponding to the computational bottlenecks of Cpufit. For details of the simulated data and fit parameters, see the Supplementary Information.

Given the parallel computing capability of the GPU, it was not surprising that Gpufit outperformed an equivalent algorithm running on the CPU. In order to verify that our code is efficiently implemented, we therefore tested Gpufit against another GPU-based fitting library: GPU-LMFit.^8^ These tests were limited to smaller datasets because GPU-LMFit is available only as a closed-source, 32-bit binary package, restricting the size of the memory it can address. Figure 3A shows the speed of the Gpufit and GPU-LMFit libraries measured as a function of the number of fits per function call (*N*), with the speed of the MINPACK library shown for reference. Both packages exhibited similar scaling in speed as the number of fits and the data size varied, however, Gpufit showed faster performance for all conditions tested. As the data size per fit was increased (Fig. 3B), the speeds became more comparable, indicating that Gpufit makes more efficient use of GPU resources for smaller fits. The increased speed comes with no loss of precision, as it was also shown that the fit results returned by Gpufit and GPU-LMFit have virtually identical numerical precision (see Supplementary Fig. S7).

**Figure 3:**
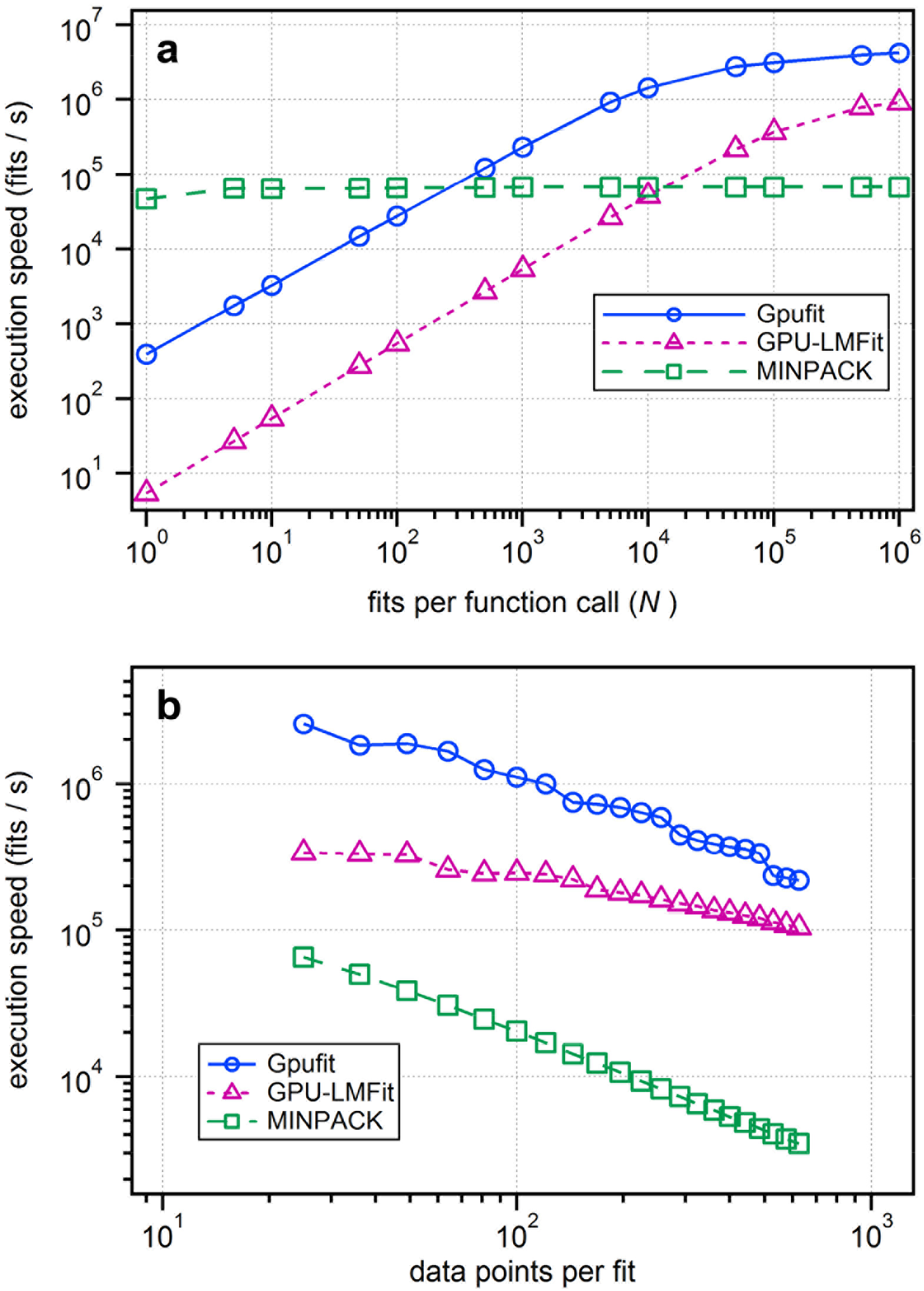
Processing speed comparison between three fitting libraries: Gpufit, and GPU-LMFit. (**a**) Execution speed vs. number of fits per function call (*N*). (**b**) Execution speed vs. number of data points per fit, where the size of the simulated 2D Gaussian peaks was varied from 5 × 5 up to 25 × 25 points. For smaller tasks (smaller numbers of fits per call or smaller data sizes) the Gpufit library makes more efficient use of GPU processing resources, outperforming GPU-LMFit by more than one order of magnitude. For larger tasks the results become more similar between the two GPU-based libraries. For details of the simulated data and fit parameters, see the Supplementary Information.

When data is subject to counting statistics (i.e. when the noise has a Poisson distribution), curve fitting using an alternative goodness-of-fit measure based on maximum likelihood estimation (MLE) has been reported to yield more accurate fit results, when compared with least-squares fitting.^9,10^ We tested the performance of this estimator using input data with simulated Poisson noise. The precision of the fit results, using either the unweighted LSE, weighted LSE, or MLE estimators, is shown in Supplementary Fig. S8. We found that the MLE estimator yielded better results than the LSE, particularly at small data values, although for larger values the two methods were approximately equivalent.

### Application to super-resolution microscopy

Gpufit is well suited for rapid processing of large datasets, and its interface allows it to serve as a drop-in replacement for existing CPU-based fit routines. To demonstrate these capabilities in a real application, we integrated Gpufit into the image analysis pipeline of a super-resolution fluorescence microscopy experiment.

Stochastic optical reconstruction microscopy (STORM) is a method for fluorescence imaging of biological samples, which obtains an image resolution significantly higher than the classical “diffraction limit” of far-field optical microscopy.^11^ This method relies heavily on image processing: the generation of a single STORM image requires thousands to millions of individual fitting operations, and this task may require minutes to hours of computation before the final image is obtained.

We integrated Gpufit into a recently published software package, Picasso,^12^ which may be used to process raw STORM data into a super-resolved image of the sample. The Picasso software is written in Python, and is optimized to carry out the fit using a multi-threaded process running on all cores of the CPU. Moreover, Picasso uses “just in time” (JIT) compilation to optimize its execution speed. We modified this section of the Picasso source code, replacing the multi-core CPU-based fitting with a call to Gpufit, as illustrated schematically in Fig. 4A. Comparing the speed of Picasso before and after the modification demonstrates the benefit of Gpufit. When analyzing a raw STORM dataset (80000 images requiring 3.6 million fit operations), a fitting task which required 99.4s with standard Picasso was completed in only 2.2s after Gpufit was included, a 45-fold speed increase, with identical fit precision. The output STORM image is shown in Fig. 4B, and a comparison of the curve fitting time with and without Gpufit is shown in Fig. 4C.

**Figure 4:**
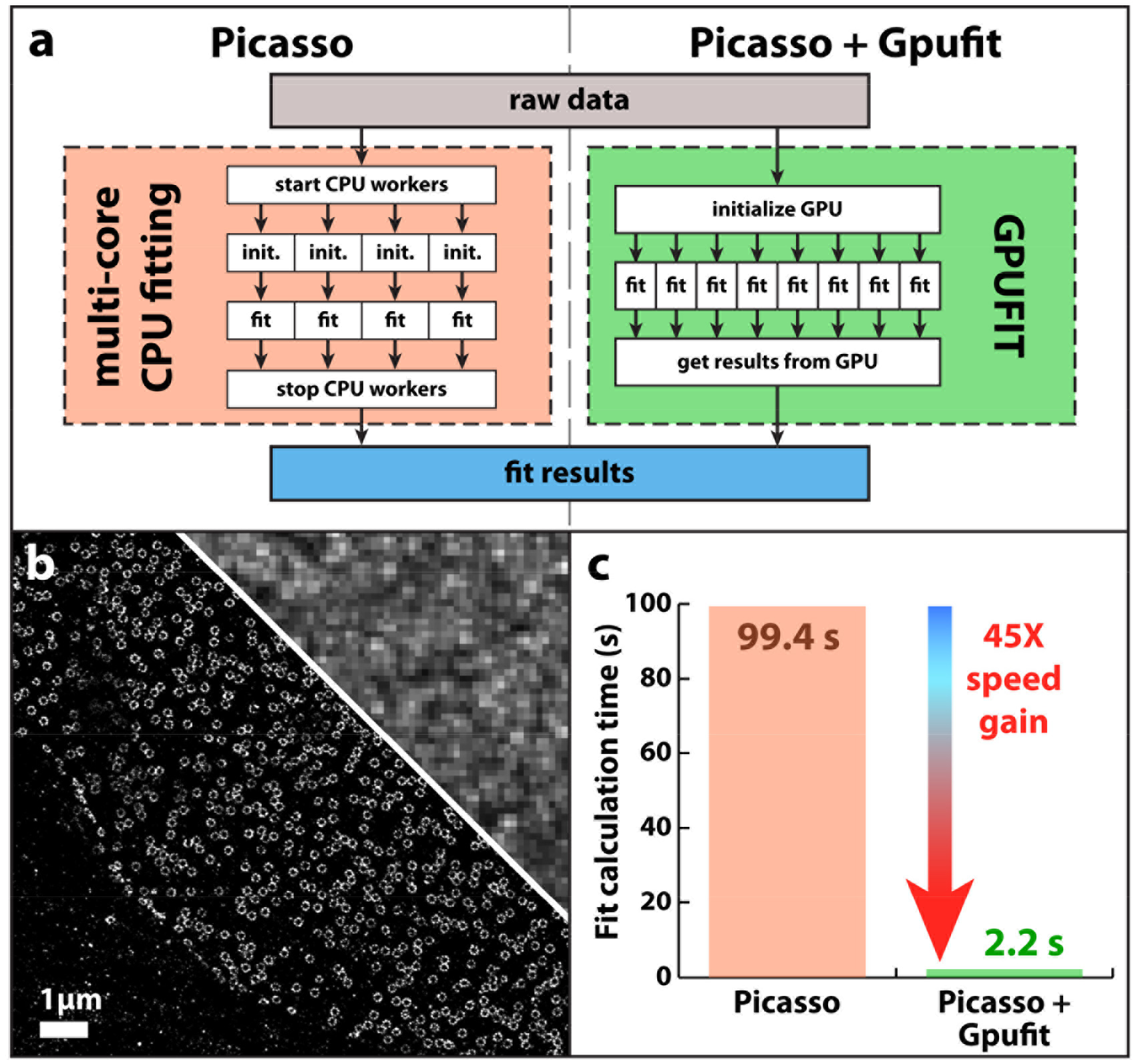
Accelerated STORM analysis in Picasso. (**a**) Schematic diagram comparing the multi-core CPU-based fitting of Picasso (left) vs. parallelized GPU-based fitting with Gpufit (right). Picasso uses multiple cores (“workers”) of the CPU to simultaneously compute the fits. We replaced this section of the Picasso source code with a call to Gpufit, which has a greater capacity for parallelization via the GPU. (**b**) Super-resolution (STORM) image of nuclear pore complexes in eukaryotic cells. The protein GP210 was labeled with a fluorescent marker (Alexa Fluor 647), and the sample was imaged on a STORM microscope (see Supplementary Information). The conventional epi-fluorescence image is shown in the top right corner, to provide a resolution comparison. Data processing was performed in Picasso, requiring 3.6 million individual curve fitting operations. (**c**) Execution time of the curve fitting process, comparing the original (published) Picasso software vs. the modified Picasso software which includes Gpufit. The processing time was reduced significantly, by a factor of 45, when Gpufit was used to handle curve fitting tasks within the application. For these measurements, the data was fit to a symmetric 2D Gaussian function (Picasso default) using the unweighted least squares estimator, and the fit size was 7 × 7 pixels. All calculations were run on an NVIDIA GeForce GTX 1080 GPU and an Intel Core i7 5820K CPU (3.3 GHz). For detailed test conditions, see the Supplementary Information.

## Discussion

General purpose GPU computing is a relatively new technology, which is making an impact in many fields of science and engineering. The software introduced here, Gpufit, represents the first general purpose, open source implementation of a non-linear curve fitting algorithm for the GPU. It is intended as a compact, high performance optimization tool, easily modified and adapted for new tasks, which can be rapidly implemented within existing data analysis applications.

In terms of performance, Gpufit exhibits similar precision and accuracy to other fitting libraries, but with significantly faster execution. In our measurements, curve fitting with Gpufit was approximately 42 times faster than the same algorithm running on the CPU. Gpufit also outperformed another GPU-based implementation of the LMA, GPU-LMFit, for all conditions tested. The absolute values of the timing results depend on the details of the fit and the computer hardware. Higher performance would be expected with a more powerful GPU (e.g. an Nvidia Tesla), or with multiple GPUs running in parallel. In addition to its speed, the principal advantages of Gpufit are its general purpose design, which may incorporate any model function or modified estimator, and the availability of the source code, which allows it to be compiled and run on multiple computing architectures.

Since Gpufit automatically distributes the curve fitting tasks over the blocks and threads of the GPU multiprocessors, the user is not required to know the specifications of the hardware, thereby allowing Gpufit to be used as a “drop-in” replacement for existing fit functions. To demonstrate this, we modified Picasso to make use of Gpufit rather than its own multi-core CPU fitting code. Despite the fact that curve fitting in Picasso was already parallelized (by virtue of its use of multi-threaded processing) we found that Gpufit outperformed the built-in Picasso curve fitting by a factor of 45 times (Fig. 4). The ease with which Gpufit was integrated into Picasso, and the gain in performance, show that our original design goals were met.

As of its initial release, the Gpufit package has several limitations. First, the fit model functions are built into the code at compilation time, and the addition or modification of a model function requires re-compilation of the source code. We also note that Gpufit requires the explicit calculation of the partial derivatives of the model, and expressions for these functions must be present in the code embodying the fit model function. However, as an open-source software project, we expect that Gpufit will continue to develop and improve, potentially removing these limitations in future versions. For example, runtime compilation of model functions written in CUDA would lift the requirement for re-building the source, and methods to approximate the derivatives numerically could also be introduced. Finally, there is the potential for porting Gpufit to other general-purpose GPU computing languages, such as OpenCL, thereby allowing the software to function on other GPU hardware platforms.

Using an inexpensive graphics card and a standard PC, we achieved speeds higher than 4.5 million fits per second for data that is typical of STORM experiments (Fig. 1). Considering recent developments in localization based super-resolution methods, in particular experiments in which >400 million individual fluorophore switching events were recorded,^13^ data processing speed has become highly relevant. A tool such as Gpufit could make the difference between a researcher waiting minutes (with Gpufit) or hours (without) before the image can be examined. This rapid feedback has a greater importance than simply reducing waiting time - it enables the quick screening of samples and test conditions, ultimately leading to higher quality image data. Furthermore, we expect that Gpufit will facilitate the adoption of new, computationally demanding data analysis approaches, such as cubic spline fitting,^14^ in order to further improve STORM image resolution.

## Conclusions

We have developed a GPU-accelerated implementation of the Levenberg Marquardt algorithm, with significantly faster performance as compared with traditional CPU-based software. Gpufit is designed to be general purpose, and as such we expect it to be useful for diverse applications in science and engineering which may depend critically on rapid data processing, e.g. high-speed feedback systems and the recently described MINFLUX method for particle tracking and nanoscale imaging.^15^ The Gpufit library is accessible from most programming environments, and the automatic configuration of GPU-specific parameters makes it simple to work with in practice. The modularity of Gpufit facilitates the addition of custom fit models or estimators, for example the mixed Gaussian/Poisson likelihood model required for accurately analyzing sCMOS camera data.^16^ Finally, Gpufit makes efficient use of GPU computing resources by exploiting simple parallelization schemes as well as distributing fit tasks evenly over the multiprocessors of the GPU to achieve high performance. Using Gpufit with our own hardware, we obtained a 42-fold improvement in execution time for batch-processing of curve fitting tasks, with no loss of precision or accuracy.

Gpufit is an open source software project, and the source code is available via the Github public software repository. Open source software development is advantageous from several standpoints: it enhances code integrity through review by users, and offers the possibility for users to fix bugs, add features, etc.^17^ In addition, open source code makes the workings of an algorithm transparent to the user, which may be a crucial factor when a “black box” software tool is not sufficient, such as in scientific data analysis applications. Finally, we note that Marquardt’s original paper introducing his algorithm, published in 1963, concludes with the following sentence: “A FORTRAN program … embodying the algorithm described in this paper is available as IBM Share Program No. 1428”.^1^ The SHARE software library referred to by Marquardt was, in fact, an early form of open source software,^18^ and we find it appropriate that Gpufit should be published in a similar manner.

## Methods

### Levenberg-Marquardt algorithm

The LMA provides a general numerical procedure for fitting a non-linear model function to a set of data points. It may be considered as a combination of the method of steepest descent and Newton’s method, having a high probability of convergence even when the initial parameter estimates are poor, and fast convergence near the minimum. The standard algorithm, as described by Marquardt,^1^ minimizes iteratively the general least squares equation:

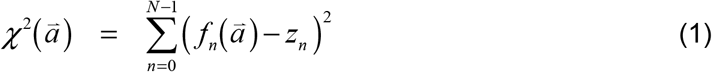

where *f*_*n*_ are the model function values, 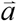 is the vector of model parameters, and *z*_*n*_ are the set of *N* data points. To find the minimum of 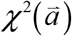 (chi-square), the algorithm performs an iterative search of the parameter space for a coordinate where the gradient of the function equals zero, i.e. 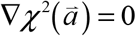. The gradient is approximated by a Taylor expansion:

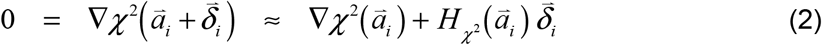

where 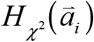 is the Hessian matrix of 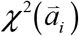, and 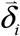 is a small correction to 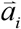 (the index *i* corresponds to the iteration number). The expression for the Hessian includes the first and the second partial derivatives of 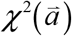, however terms containing the second derivatives are assumed negligible and ignored. Solving (2) for 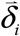 yields the Newton step for the minimization of 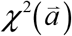. Up to this point, the LMA is equivalent to Newton’s Method.

A special characteristic of the LMA is the damping factor *λ* which controls the step size of each iteration by modifying the diagonal elements of the Hessian:

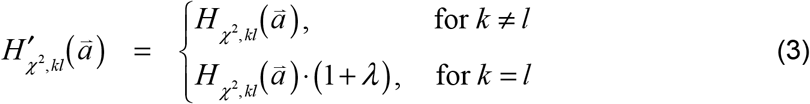

where *k* and *l* are the matrix indices. The positive factor *λ* is initialized with a small value, and thus initially the algorithm behaves like Newton’s method. As the algorithm iterates, if the value of chi-square in the latest iteration is smaller than in the previous step, *λ* is decreased by a constant factor *v*. Otherwise, *λ* is increased by the same factor. Increasing *λ* causes the LMA to tend towards the behavior of the method of steepest descent. In this manner, the LMA adjusts between the two methods, as the minimum is approached.

By transposing equation (2) and applying the damping factor (3), a system of linear equations is obtained which may be solved for 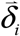, e.g. by the Gauss-Jordan method:^3^

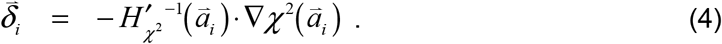

If the iteration is successful (chi-square decreased), the difference 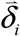 is added to the previous parameter values:

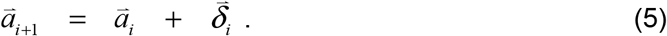

The damping factor *λ* is updated after each iteration, and convergence is tested. Any convenient convergence criterion may be used, but in general the overall convergence of the LMA depends on the relative size of the parameter adjustment in each iteration. As originally set out by Marquardt, with a reasonable choice of *r* and *ε* (e.g. *r* = 10^−3^ and *ε* = 10^−5^), the algorithm has converged when

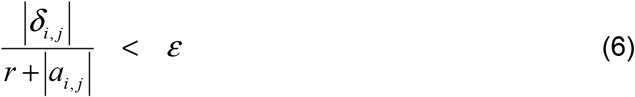

is satisfied for all parameters, where *r* is a small positive constant (to avoid division by zero), *i* is the iteration number, and *j* is the parameter index.

### Estimators of best fit

Least squares estimation (LSE) is a common method for finding the parameters which yield the minimal deviation between observed data and a model function. The standard LMA minimizes the general LSE formula given by equation (1). However, it is also possible to include weighting factors in the calculation of chi-square, for example:

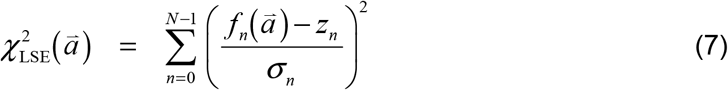

where *σ*_*n*_ represents the uncertainty (standard deviation) of the data. This allows the precision of each data point to be taken into account.

In cases where the uncertainties of the data points are Poisson distributed, a maximum likelihood estimator (MLE) yields more precise parameter estimates than the LSE.^9,10^ In this situation it is beneficial to use an alternative estimator with the LMA. A procedure has been described^6^ in which the LSE formula (1) is replaced by the MLE equation for Poisson deviates, as follows:

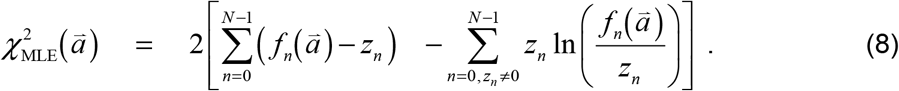

Using this estimator within the context of the LMA is relatively simple to implement, requiring only the calculation of the gradient and Hessian matrix of 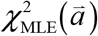, and the calculation of 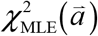 itself. As before, in the calculation of the Hessian matrix, terms containing second partial derivatives are ignored.

### Gauss-Jordan elimination

Gauss-Jordan elimination was used to solve the system of equations (4) in the LMA, and a parallelized Gauss-Jordan algorithm was developed for this purpose. The algorithm was parallelized according to the elements of the augmented matrix, with each thread responsible for one element of the matrix as the left side is transformed to reduced row echelon form. Partial pivoting (row swapping) was used to ensure precise and numerically stable calculations.^3^ The sorting step in the pivot operation was accomplished on the GPU by means of a parallel bitonic merge sort.^19^ The details of this algorithm are fully documented in the Gpufit source code.

### Software for comparison tests

To evaluate its performance, Gpufit was compared against an equivalent CPU-based algorithm (Cpufit) and two other curve fitting libraries: MINPACK^7^ and GPU-LMFit.^8^ Cpufit is a standard implementation of the LMA based on published examples,^1,3^ which we wrote in C++ for execution on the CPU. C++ Minpack is an open-source C/C++ implementation of MINPACK which runs on the CPU (we used the function `lmder()` from this library).^7,20^ Both C++ Minpack and Cpufit were run in single CPU threads. GPU-LMFit is a closed-source implementation of the LMA (32-bit binary files are publicly available) which runs on the GPU and provides the option of using LSE or MLE as the estimator.^8^

### Computer hardware

All tests were executed on a PC running the Windows 7 64-bit operating system and CUDA toolkit version 8.0. The PC hardware included an Intel Core i7 5820K CPU, running at 3.3 GHz, and 64 GB of RAM. The graphics card was an NVIDIA GeForce GTX 1080 GPU with 8 GB of GDDR5X memory. All source codes, including the Cpufit, Gpufit, and C/C++ Minpack libraries, were compiled using Microsoft Visual Studio 2013, with compiler optimizations enabled (release mode). Additional details of the test conditions and instructions for compiling the Cpufit and Gpufit software libraries are provided in the Supplementary Information and the Gpufit documentation.

### Source code availability

The source code for Gpufit, including external bindings, is available for download from a public software repository located at http://www.github.com/gpufit/Gpufit. The source code for the CPU-based reference algorithm, Cpufit, is also included in this repository. The Gpufit User’s Manual, describing how to use the software and containing instructions for building and modifying the source code, may be found online at http://gpufit.readthedocs.io.

### Data availability

The datasets generated during and/or analyzed during the current study are available from the corresponding author on reasonable request.

## Acknowledgements

We would like to thank Dr. C. Wurm and Dr. E. Rothermel for the preparation of the sample used for STORM imaging, and also Dr. V. Cordes for providing the primary antibody against GP210. We thank N. Warmbold for contributions to an early version of the Gauss-Jordan algorithm used in the Gpufit source code. We thank Dr. M. Roose for fast and effective IT support. We thank Prof. Dr. S. W. Hell for generous support in the form of funding and equipment. M.B. gratefully acknowledges funding from the European Molecular Biology Organization (ALTF 800-2010) and the Max Planck Society.

## Author Contributions

M.B., B.T., and B.S. conceived the project. A.P., B.T., J.K., and M.B. wrote the Cpufit and Gpufit source code. A.P. and M.B. carried out the quantitative evaluation of Gpufit. M.B. performed the STORM experiment. A.P. and J.K. wrote the external bindings for Gpufit. J.K. created the usage examples. J.K., A.P., M.B., and B.T. wrote the documentation. M.B., B.T., and B.S. supervised the research. M.B. wrote the manuscript.

